# Sequence learning attenuates cortical responses in both frontal and perceptual cortices in early infancy

**DOI:** 10.1101/2021.11.10.468062

**Authors:** Sagi Jaffe-Dax, Anna Herbolzheimer, Vikranth Rao Bejjanki, Lauren L. Emberson

## Abstract

Prior work has found that the frontal lobe is involved in higher-order sequential and statistical learning in young infants. Separate lines of work have found evidence of modulation of posterior sensory cortices during and after learning tasks. How do these processes relate? Here, we build evidence the infant frontal lobe was modulated during sequential learning and ask whether posterior perceptual cortices show corresponding modulation. First, replicating and extending past work, we found evidence of frontal lobe involvement in this task. Second, consistent with our hypotheses, we found that there is a corresponding attenuation of neural responses in the posterior perceptual cortices (temporal and occipital) to predictable compared to unpredictable audiovisual sequences. This study provides convergent evidence that the frontal lobe is crucial for higher-level learning in young infants but that it likely works as part of a large, distributed network of regions to modulate infant neural responses during learning. Overall, this work challenges the view that the infant brain is not dynamic and disconnected, lacking in long-range neural connections. Instead, this paper reveals patterns of a highly dynamic and interconnected infant brain that change rapidly as a result of new, learnable experiences.

## Introduction

Extracting information from recurring patterns in the environment is a crucial ability that underlies infant learning across numerous domains (Saffran and Kirkham, 2018). From inferring meaningful information out of sensory input through developing complex cognitive skills, humans rely on detecting re-occurring patterns and identifying pattern-violations starting early in life (Arnon, 2019; Newman et al., 2006). Broadly, we consider these abilities to arise from statistical learning. Recurring patterns manifest as statistical information in sensory input. Humans are highly sensitive to statistical information and learn from statistical information incidentally starting early in infancy (need citations). While behavioral studies have made significant progress in understanding the computational mechanisms that underlie statistical learning (Siegelman et al., 2019; Thiessen, 2017), the underlying neural mechanisms that support statistical learning are not yet well understood (Schapiro et al., 2012; Turk-Browne et al., 2009) and there is even less known about the brain regions involved in statistical learning in infants (though see Ellis et al., 2021; Emberson et al., 2015; Kabdebon et al., 2015; Kersey and Emberson, 2017). Importantly, understanding the neural mechanisms supporting statistical learning in infancy will shed light on the mechanisms supporting development, and provide a foundation upon which we can investigate how these learning mechanisms change across conditions, stimuli and over development. Here, we use functional near-infrared spectroscopy (fNIRS) to investigate whether the predictability of a recurring sequence can result in top-down attenuation of sensory cortices and whether the frontal lobe is a potential source of this top-down information.

We focus our current investigation on the difference in neural processing of predictable vs. unpredictable sequences of audiovisual stimuli. Classic statistical learning paradigms have segments of predictability (e.g., within a word) and unpredictability (e.g., across words, Saffran et al., 1996). Given the slow temporal resolution of fNIRS, we distilled this key characteristic of statistical learning into separate sequences: one of which is entirely predictable and the other is much less predictable. We can then contrast neural responses to predictive vs. unpredictable sequences to elucidate how the infant brain changes when an infant is able to learn and predict from recurring patterns in the environment.

It has long been shown that already at early infancy, recurring sequences can be learned (Marcus et al., 1999; Saffran et al., 1996). As early as immediately after birth, neonates respond selectively to patterned stimuli (Benavides-Varela and Gervain, 2016; Gervain et al., 2008). Starting at 2 months of age, infants were already able to show behaviorally that they learned sequences of two consecutive events (Kirkham et al., 2002; Slone and Johnson, 2016), or co-occurrences of two items in space (Fiser and Aslin, 2002). It has been behaviorally demonstrated that babies can learn sequences of three items such as in the context of rule-learning (Ferguson et al., 2018; Johnson et al., 2009; Lew-williams et al., 2017; Saffran et al., 2007; Slone and Johnson, 2018). Sequence learning, as a type of statistical learning, is believed to be important for language acquisition (Gomez and Gerken, 1999; Lew-Williams and Saffran, 2012; Pelucchi et al., 2009; Potter and Lew-Williams, 2019). With one exception for infants over 1 year old (Schonberg et al., 2018), there are no behavioral studies that have demonstrated learning of 4 item sequence in infants. While there is a good amount of behavioral evidence that infants engage in learning of recurring and predictable sequences of stimuli, little is known about the neural mechanisms supporting this ability.

However, a recent EEG study investigated sequence learning of 4 items in 3-month-old infants and implicated the frontal lobe in this process. Specifically, Basirat et al. (2014) found that in early infancy the left frontal lobe attenuates its response to deviant stimulus when it is more predictable based on consistent serial sequence of stimuli. The researchers presented a sequence of trials (block) consisting of mostly four identical stimuli (XXXX) interleaved with rare trials which had a deviant stimulus at the end (XXXY) to a group of infants. They compared their responses to the deviant with another block of trials where the common trial contained a deviant (XXXY) and the rare trials had no deviant (XXXX). This design allowed them to disentangle low-level effects like repetition suppression from more high-level effect like predictability and top-down modulation arising from predictability. Their findings show that expecting a deviant stimulus (in the second block) attenuated the deviant-elicited response in a selection of electrodes on the left frontal scalp compared to when infants did not expect the deviant (the first block). Thus, there is an attenuation of neural responses to stimuli when they are expected or predictable. This finding dovetails with other work demonstrating that the infant frontal lobe is involved in processing of familiarity and novelty (Nakano et al., 2008) and is involved in rule learning (Gervain et al., 2008; Werchan et al., 2016) and learning audiovisual associations (Kersey and Emberson, 2017) starting early in life, and broader proposals that the infant frontal lobe is available to contribute to infant cognition and learning starting early in life (Dehaene-Lambertz and Spelke, 2015; Grossmann et al., 2013). Thus, a recent study has found that the frontal lobe appears to be involved in learning a sequence of 4 stimuli.

The current study will replicate and extend the finding that the frontal lobe is involved in learning predictable sequences in early infancy using fNIRS. One of the major benefits of fNIRS is the superior spatial resolution of this neuroimaging method as source-localization is not needed. To ensure accuracy in our spatial localization, we co-register the probe localizations with MR templates using a video-based method to allow anatomical localization of fNIRS recordings with infants (Jaffe-dax et al., 2020). Moreover, we define our regions of interest *a priori* based on the localization of each probe for each infant and the underlying neuroanatomy. In addition, we employ non-speech, novel audiovisual stimuli (see Figure 1). Thus, we will extend the findings from Basirat et al (2014) to different stimuli and using neuroimaging modality with better spatial resolution in order to provide stronger evidence that the frontal lobe is involved in sequence learning in young infants.

**Figure 1.**
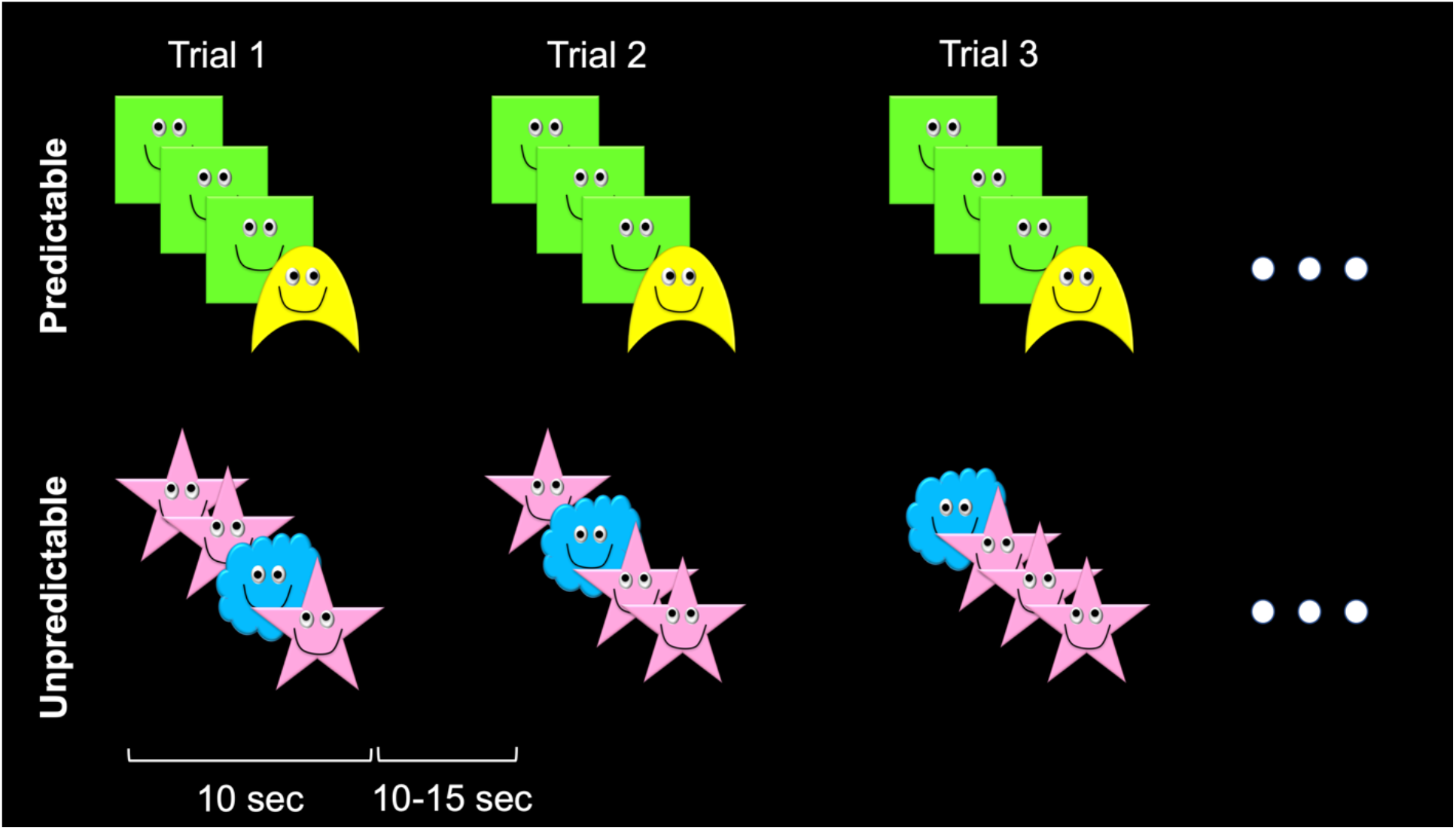
Stimuli and experiment structure. The experiment consisted of two conditions: Predictable and Unpredictable. Each condition consisted of 8 consecutive trials. In each trial, four shapes moved sequentially across the screen accompanied with a unique sound. In the Predictable condition, the order of the shapes was consistent between trials (AAAB). In the Unpredictable condition, the order of the shapes varied between trials (AABA, ABAA, BAAA, but not AAAB).

In addition, our extension of Basirat et al (2014) will determine whether learning the predictability of a sequence results in top-down modulation of posterior sensory regions. Based on the framework of predictive coding (Friston, 2005; Rao and Ballard, 1999), which posits that neural signals in the cortex communicate feedback/top-down prediction about upcoming sensory input and feed-forward/bottom-up signals of prediction error (arising from the comparison of sensory input to predictions), we hypothesized that learning of predictable sequences would be associated with reduced prediction error and thus **attenuated response in both associative and in sensory cortices**. Previous work in adults found that repetition of presented stimuli attenuated neural responses specifically when the repetition was expected, suggesting that the source of repetition-induced suppression reside in the predictability of the presented stimuli (Summerfield et al., 2008). On the other hand, the response to a deviant stimulus embedded in a sequence of standard stimuli were also attenuated by predictability in adults (Sussman et al., 2003). Recently, the impact of predictability was demonstrated in young infants, where unexpected violation of learned association augmented neural responses (Emberson et al., 2015; Kouider et al., 2015) in sensory cortices. We now set out to explore the role of this prediction-induced process in statistical learning of longer time-scales. Namely, we aim to evaluate whether the emergence of predictive processing in sensory cortices is part of the neural mechanisms associated with statistical learning in infancy.

We recorded cortical activity of infants’ Frontal lobes and two posterior sensory cortices (Temporal and Occipital lobes) in two experimental conditions: 1) Predictable condition, where in each trial, four shapes were presented in a consistent order of AAAB (three standard stimuli followed by a deviant). 2) Unpredictable condition, where the order of the shapes differed from trial to trial (AABA, ABAA or BAAA). We chose these trial structures across the two conditions to differentiate between two alternatives. If the consistency of the shapes order between trials was not learned, and the magnitude of the cortical response was only governed by the appearance of the deviant shape in each condition, then we would expect the Predictable condition with the AAAB pattern to yield a stronger response in posterior sensory cortices; the deviant appears after three consecutive standards compared to zero, one or two standards in the unpredictable condition (Yaron et al., 2012). If, on the other hand, as we hypothesized, infants were able to learn the consistency of sequence, use this learning to predict upcoming sensory input and communicate this prediction through top-down or feedback connections to posterior cortices, we will observe that the Unpredictable condition would yield a stronger cortical response in posterior cortices. Specifically, the stronger cortical response in the Unpredictable condition reflects the relative attenuation of cortical responses to the (predicted) deviant stimulus in the Predictable condition and a strong cortical response to the (unpredictable) deviant in the Unpredictable condition. Thus, the specific structure of the trials allowed us to make inferences about how the temporal predictability of the deviant stimulus affects perceptual despite not having the temporal resolution to investigate responses to individual stimuli (as in EEG). Indeed, predictable deviants elicited weaker deviant-related response in kids and adults (Max et al., 2015; Sussman et al., 2003) and in the frontal lobe in Basirat et al (2014). We hypothesize to find the same attenuation of deviant responses in the posterior perceptual systems of young infants.

Compared to other studies investigating responses to odd-ball stimuli, the current study design allows us to distinguish between simple deviance detection (i.e., that X is presented more frequently and thus seeing Y evokes a strong perceptual response), that can be processed locally, and more complex responses to sequential regularities that would require the involvement of higher-level cortices like the frontal cortex and feed-back or top-down connections from higher level cortices to the posterior perceptual cortices.

## Methods

### Participants

Twenty-seven infants at 5-7 month of age were included in the analysis. The experiment was terminated after both conditions were recorded or when the infant looked away from the screen for more than half of a trial duration. Ten additional infants were excluded from the analysis for not completing at least 5 trials of each condition. The study was approved by the university’s Institutional Review Board and informed consent was obtained before the beginning of the study from a legal guardian of the infant. Families received $10, a t-shirt and a children’s book for their participation.

### Stimuli

Each trial consisted of four shapes crossing the screen sequentially while an associated sound was played. The shapes consisted of green square, yellow crescent, pink star, and blue cloud. Each shape was added caricature eyes and smile (Fig. 1). The shapes crossed the screen in one of four directions: up, down, left, or right. During the movement of the shapes on the screen, a sound was played. The sounds were old windows startup, rattle, train, or chimes. Each trial lasted for 10 seconds and the Inter-trial intervals were jittered between 10 to 15 seconds during which time a dimmed fireworks video and nursery song was played to keep infant attention towards the screen (Emberson et al., 2015). Each condition consisted of eight consecutive trials. In the Predictable condition, the order of the shapes within each trial was consistently AAAB (Fig. 1 top). In the Unpredictable condition, the order of the shapes within each trial was pseudo-randomly chosen between AABA, ABAA, and BAAA (and not AAAB; Fig. 1 bottom). The associations of shapes, movement direction, sounds and conditions were pseudo-randomly assigned for each infant. The order of conditions was counter-balances between infants.

### Procedure

Infants sat on their caregiver’s laps in front of a screen at approx. 60 cm distance from the screen. Stimuli moved across the central part of the screen for about 20 degrees of visual angle. Associated sounds were played through central positioned speakers at a comfortable level. The fNIRS cap was placed on the infant’s head, infrared sensors sensitivity was adjusted and photogrammetry measurements were taken, while infant’s attention was distracted with nursery rhymes.

### fNIRS recording and analysis

We recorded infants’ cortical activity from 74 fNIRS channels at ~13.33 Hz (inter-sample interval of 75 ms) at three wavelengths (790, 805 and 830 nm) using custom-made infant fibers, connected to Shimadzu LABNIRS. Channel separation on the scalp was 25 mm. Cap position on the infant’s head was estimated using photogrammetry of fiducial points (Nz, Cz, Iz, AL, AR and cap edges; Lloyd-Fox et al., 2014). Channel position was interpolated and projected to MNI space using SPM-fNIRS (Tak et al., 2016). Each channel was assigned to its containing lobe. For each lobe, we averaged the channels that were assigned to it, so that the signal that was recorded from each lobe could be averaged across infants.

Preprocessing was performed for each channel using Homer2 toolbox for Matlab (Huppert et al., 2009) with the recommended parameters from (Brigadoi et al., 2014). Raw intensity data was converted to optical density. Motion artifacts were detected using tMotion = 1, tMask = 1, STDEVthresh = 50, and AMPthresh = .5. Epochs containing detected artifacts were removed from the data and spline interpolated using p = 0.99. Data was then band-pass filtered between 0.01 and 1 Hz. Filtered optical density from the three wave lengths was transformed to HbO_2_, HHb and total Hb concentration changes using modified Beer-Lambert equation with partial path factor (ppf) of 6 mm for all three wavelengths. Any channel containing concentration changes beyond ±5e-5 μM were omitted from further analysis. Pre-processed data was averaged within each lobe and homologous lobes were aggregated together. We recorded from 17 to 22 channels in the Frontal lobes; from 29 to 40 channels in the Occipital lobes; and from 13 to 22 channels in the Temporal lobes. The first trial of each condition was omitted from analysis. We averaged the response for each condition from 2 seconds before the onset of the first shape in the trial to 8 seconds after the offset of the fourth shape (epoch length of 20 seconds). We also assessed whether there were changes in task-based functional connectivity, between the conditions, using a background connectivity approach (Al-Aidroos et al., 2012). More details on these methods as well as our results are included in the Supplementary Materials.

## Results

For each lobe, we used Monte-Carlo cluster size correction (Maris and Oostenveld, 2007) to calculate the probability of finding a significant cluster of (temporally consecutive) time samples for which there was a significant difference between the conditions. We found a stronger response for the Unpredictable condition compare to the Predictable condition in the Frontal, Occipital and Temporal lobes. In the Frontal lobes, a significant difference between conditions was found between 12.45 – 14.4 sec post trial onset (Fig. 2 top; cluster *P* < 0.05). In the Occipital lobes, we found a significant difference between conditions between 10.35 – 13.5 sec post trial onset (Fig. 2 middle; cluster *P* < 0.05). In the Temporal lobes, we found a significant difference between conditions between 12.6 – 14.18 sec post trial onset (Fig. 2 bottom; cluster *P* < 0.05).

**Figure 2.**
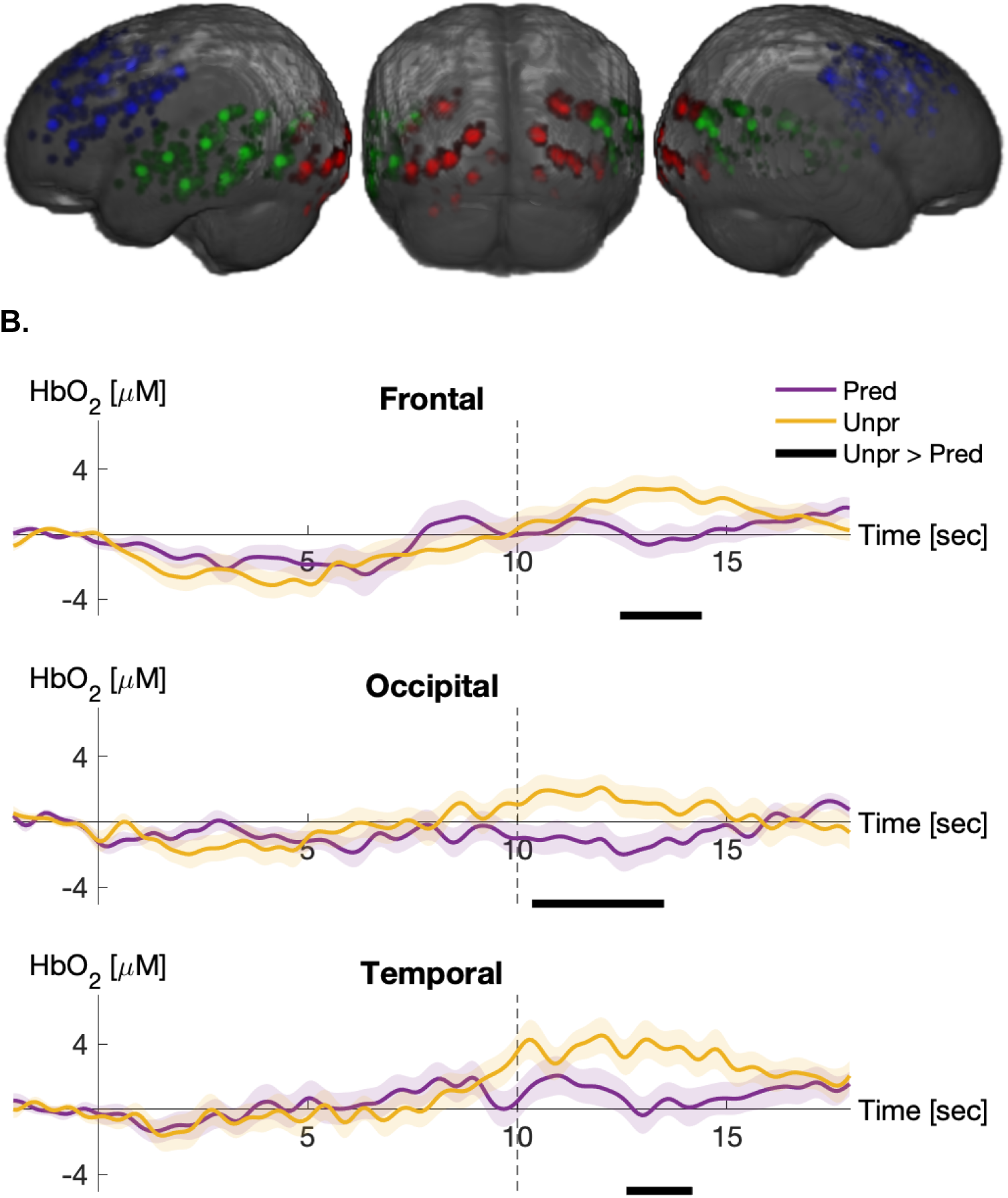
**A.** Spatial co-registration of the fNIRS channels by lobe. Each patch denote a single channel in an individual participant. Overlapping locations are represented in saturation from pale to vibrant color. Blue patches – channels on the frontal lobe; green – temporal; red – occipital. **B.** fNIRS response to the Predictable (purple) and the Unpredictable (orange) conditions by lobe. Concentration changes of oxyhemoglobin (HbO_2_) as a function of time in seconds from trial onset. Stimulation offset is denoted by the dashed line at 10 seconds from trial onset. Shaded patches represent SEM. Black bars under each graph denote the time windows where the response for the Unpredictable condition was greater than the response for the Predictable condition with cluster-p < 0.05.

In addition, we compared the concentration changes in deoxyhemoglobin (HHb) between conditions and did not find any time window of significant difference (Fig. S1). These results verified that the condition difference that we found in the HbO_2_ did not stem from difference in the overall blood pressure between conditions.

We hypothesized that learning the consistent pattern of shapes in the Predictable condition would parallel attenuation of cortical response in the lobes that were measured. We estimated the trajectory of change in response amplitude as a function of trial number within each condition to track the dynamics of learning in the Predictable condition. For each trial, we averaged the HbO_2_ response from each lobe in the respective time window post trial onset as found in the previous result for each lobe separately (Fig. 2). Figure 3 depicts the trial-by-trial change of HbO_2_ concentration by condition. In the Temporal lobe, we found a significant interaction between condition and trial number (*F*_7,397_ = 2.15, *P* < 0.05). The interactions in the Frontal and Occipital lobes did not reach significance (*F*s = 1.05 and 1.47 respectively, *P*s > 0.1). These findings suggest that learning the consistent pattern of shapes in the Predictable condition was associated with reduction in activity in the Temporal lobe.

**Figure 3.**
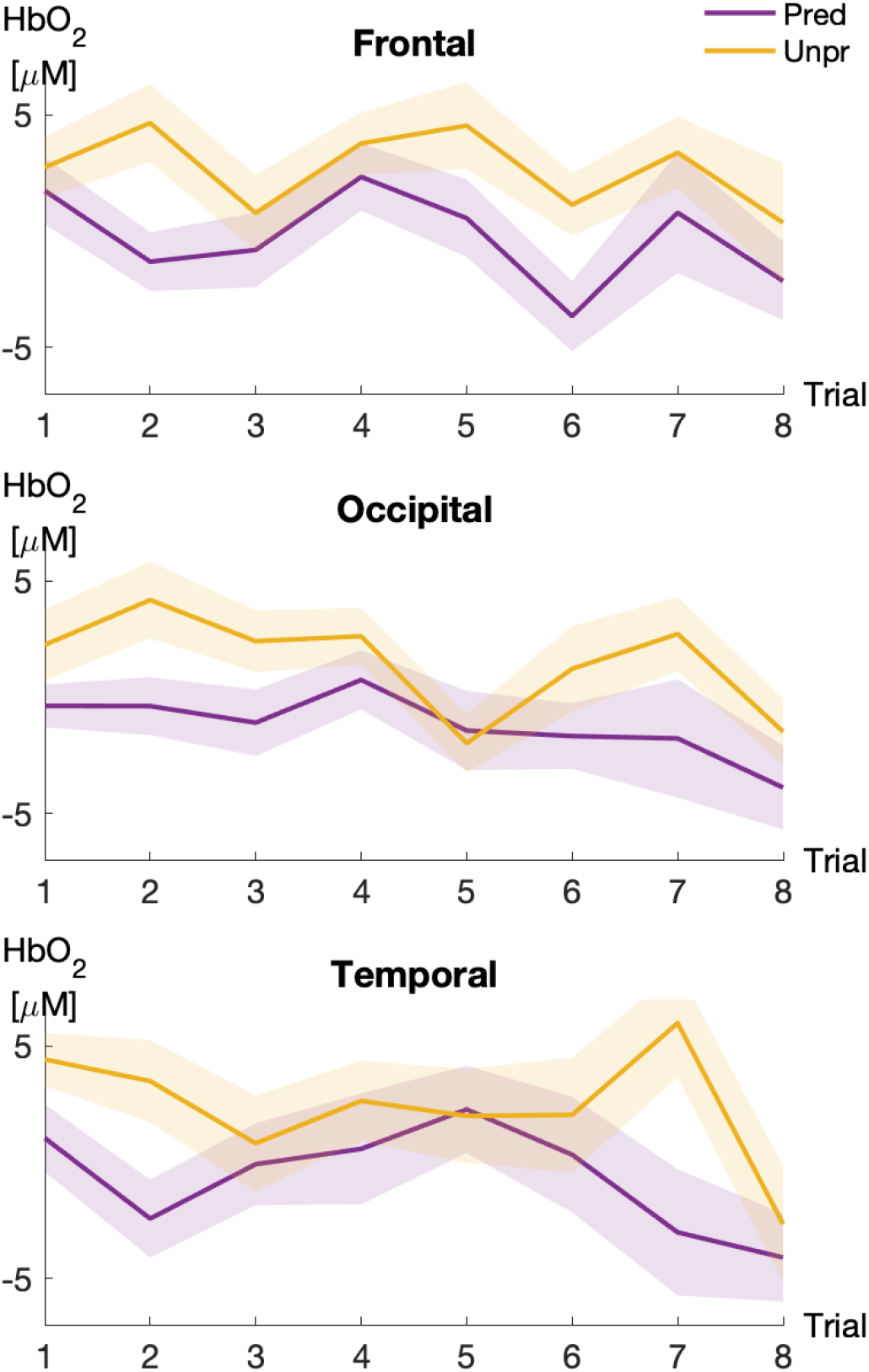
Trial-by-trial fNIRS amplitude change by condition. Mean HbO_2_ concentration change as a function of trial number for Predictable (purple) and unpredictable (orange) conditions. Shaded patches represent SEM.

We also hypothesized that we would find differences in task-based functional connectivity between the two conditions. Specifically, top-down signals from the frontal lobe are a likely mechanism underlying the modified responses to sensory input observed in the Predictable condition. Thus, we hypothesized that there would be greater connectivity between the frontal lobe and posterior sensory regions in the Predictable rather than the Unpredictable condition. To test this hypothesis, we developed a pipeline to assess task-based functional connectivity, using a “background connectivity” approach, with infant fNIRS data (see Supplementary Materials for further details including a systematic evaluation of some key analytic decisions in this pipeline). This pipeline did not reveal any reliable differences in task-based connectivity, between frontal, occipital and temporal cortices, between the two conditions. We discuss possible reasons for this in the Discussion.

## Discussion

Using fNIRS, we recorded neural activity in Frontal, Temporal and Occipital lobes in 6-month-old infants while infants engaged in a sequential audiovisual learning paradigm (Predictable condition) and a frequency-balanced control condition (Unpredictable condition, within subjects, condition order counterbalanced). We found that the effect of predictability is reflected in every recorded region. This pattern of results provides evidence for a) the frontal lobe involvement in sequential learning and replicates and extends previous work with younger infants using EEG using a very similar paradigm (Basirat et al., 2014) and adds to broader evidence that the frontal lobe is involved in various types of learning in infancy including statistical learning (Ellis et al., 2021), audiovisual associative learning (Emberson et al., 2015; Kersey and Emberson, 2017; Kouider et al., 2015); b) provides from suggestive evidence that these learning based changes are not constrained to the frontal lobe but are also present in posterior perceptual cortices suggesting a role for the frontal lobe and feedback neural connections in these posterior changes; c) that the infant brain is highly plastic and readily shaped through experience with patterns and relatedly learning.

The first major finding is a further solidified link between the frontal lobe in infancy and learning of sequential or higher-order statistical properties of the sensory input. The two conditions in this experiment differ not in the low-level statistical properties of the stimuli (e.g., how frequent individual stimuli are) or the stimuli themselves (all stimuli were randomized across conditions), but in the higher-level statistical properties of the stimuli or the sequences in which the stimuli occurred. Thus, detecting the differences between the Predictable condition and the Unpredictable condition in our paradigm requires integration of information across long timescales both within a given block as well as across repetitions of the Predictable blocks as this pattern would only be detectable across repetitions of these sequences. The infant frontal lobe has been implicated in processing stimulus novelty (Nakano et al., 2009), in rule learning (Gervain et al., 2008; Werchan et al., 2016) as well as in learning audiovisual associations (Kersey and Emberson, 2017). This study builds on a particular demonstration that the frontal lobe might be specifically involved in processing stimuli in sequential contexts: Basirat et al. (2014) used EEG to determine that in early infancy the left frontal lobe attenuates its response to deviant stimulus when it is more predictable based on consistent serial sequence of stimuli. The current experiment extends this finding from speech to non-speech audiovisual stimuli, from EEG to fNIRS (which provides greater spatial resolution and certainty compared to EEG) and to a slightly older age of infant (4 to 6 month olds). We find a broadly consistent pattern with Basirat et al (2014): in conditions where deviant responses are predictable in a sequential context, responses in the frontal lobe are attenuated. One difference between the studies is our finding of bilateral attenuation of neural responses as opposed to the response being localized to the left frontal lobe as reported in Basirat et al (2014). This difference may be attributable to the difference in stimuli as Basirat et al (2014) employed speech stimuli which may be subject to left hemisphere specialization by this stage of development. Thus, our findings replicate and extend the previous finding from Basirat et al (2014) and broadly support proposals that the infant frontal lobe is available to support infant perception and cognition (Dehaene-Lambertz and Spelke, 2015; Grossmann et al., 2013). We find that the frontal lobe may be particularly involved in the contextual processing of stimuli and higher-order learning.

The second major finding of this paper is that the impact of sequential learning and stimulus predictability is not constrained to the frontal lobe but is also present in posterior sensory cortices. Consistent with our hypotheses, we found an attenuation of neural responses when infants are able to learn the higher-order sequence and consequently predict the upcoming audiovisual stimuli. We infer that the attenuated response to Predictable condition compared to Unpredictable condition reflects a reciprocal process between posterior perceptual cortices (i.e., the occipital and temporal lobes) and higher-level, associative cortices (i.e., frontal lobe). In particular, this mechanism would implicate feedback neural connections and is broadly consistent with the Predictive Coding framework (Friston, 2005; Rao and Ballard, 1999), where cortical activity reflects prediction error, or deviances from predictable input. As the pattern of events is learned and events become more predictable, their processing elicits weaker cortical responses. Crucially, this finding has important implications for the view that feedback neural connections are available early in life and are not subject to a protracted developmental trajectory (Amso and Scerif, 2015). This also provides an important avenue through which perceptual cortices can be modulated based on experience in the world which is distinct from both the two dominant theories arguing for either bottom-up, passive absorption of sensory input or maturational/critical period constraints (Maurer and Werker, 2014).

The current finding that prediction attenuates neural responses in perceptual cortices substantially extends previous work showing that the occipital lobe can be modulated based on feedback neural connections (Emberson et al., 2015). First, the previous finding only demonstrated modulation of the occipital lobe but the current finding provides additional evidence that the temporal lobe can also be modulated through feedback neural connections. Second, the current finding extends both the complexity of the learning context (i.e., expanding from a simple audiovisual association to processing the relative predictability of a deviant stimulus based on sequential context or higher-order statistical properties of the input). This change in learning context where perceptual cortex modulation occurs is significant because a) it directly implicates a specific higher-level cortex (i.e., the frontal lobe) in the learning and provides evidence for the origin of the modulation for sensory cortices and b) extends the contexts in which the infant brain will employ feedback to modulate posterior cortices based on predictability both in terms of the length of the feedback connections and the complexity of the task. These findings also extend Jaffe-Dax et al., (2020), where trial-by-trial prediction error was tracked in 6-months-old infants using a combination of fNIRS and computational learning models in the identical task to Emberson et al. (2015). This approach revealed that prediction error is processed in frontal lobe and propagated back towards posterior cortices. These findings are highly consistent implicating the frontal lobe in the modulation of perceptual cortices and particularly in the context of processing higher-order statistical information and information that occurs over longer time-scales.

Finally, this work broadly suggests that, starting early in life, the infant brain is highly dynamic, interconnected and plastic. We find that the infant brain is able to modulate neural activity in response to new and highly complex stimuli within a matter of minutes and likely uses a combination of feed-forward and feedback neural connections to do so. This is striking given that there is substantial evidence that the infant brain increases in long-range connectivity both functionally (Bulgarelli et al., 2020; Gao et al., 2017; Homae et al., 2010) and structurally (Dean et al., 2015) throughout the first two years of postnatal life. Our findings therefore support the notion that young infants are able to employ these nascent connections to form large-scale neural networks and thus modulate neural activity broadly in response to new experiences.

However, a major limitation of the current paper is that direct analytic investigations into the background connectivity patterns in this task did not reveal differential connectivity patterns across the two conditions. It is important to note that the application of these analytic procedures to infant fNIRS is novel and the understanding of how to best reveal task-based functional connectivity will likely evolve substantially in the coming years. To this end, we present the pipeline we utilized, and evaluate the influence of some key analytic decisions, in the Supplementary Materials. There are several likely reasons for not finding differences in connectivity and a number of future avenues to explore. First, even though this study employs many more fNIRS channels than are typically used in an infant fNIRS study, fNIRS still has a much more restricted field of view than fMRI. Thus, it is quite possible that there are regions and networks involved that we were not able to record from. For instance, we did not focus our data collection on the parietal lobe but did have a small number of channels localized to that region in our infants, and exploratory analyses of these regions suggests that the parietal lobe might be differentially engaged (along with the frontal lobe) in this task. However, it is also worth noting that there are key neural regions that cannot be recorded from with contemporary fNIRS systems. For example, in both infants (Ellis et al., 2021) and adults (Turk-Browne et al., 2009), it was reported that the hippocampus played a role in supporting learning of serial dependency, and specifically in predicting upcoming events (Kok and Turk-Browne, 2018). Relatedly, it is likely that feedback connections are not specifically available to the frontal lobe and under other task conditions other cortical regions could be involved in modulating infant perception. For example, behavioral work by Xiao and Emberson (2019) argued that the amygdala was modulating regions involved in face perception by 9 months of age. Neuroscientific investigations of other regions and other tasks would determine the generality vs. specificity of these findings.

Another important point of consideration is that it is difficult to dissociate prediction from other cognitive processes including attention and learning. In particular, given that attention is modulated by predictability as well as learning, it is likely that attention varies across these task conditions though it is not clear in what direction. We controlled for attention in terms of monitoring looking time and task compliance online during the task with no differences in numbers of blocks completed etc across conditions. Thus, at a low-level (i.e., looking at the stimuli) there are not differences in overt attention across conditions. However, it is likely that the depth of attention differs both within a condition (e.g., early vs. late) as well as between conditions. The more general question of how prediction is related to attention is one that is of active investigation in the mature brain (e.g., Summerfield and de Lange, 2014). At present, this work requires highly demanding tasks that are beyond the scope of what is possible in infants (e.g., Summerfield and Egner, 2016)

In sum, we employed fNIRS to investigate how sequential predictability modulated responses not only in a region hypothesed to be essential for learning these sequences (i.e., the frontal lobe) but also in regions where feedback connections can modulate perceptual processing based on predictability (i.e., the occipital and temporal lobes). We find convergent evidence with previous work that the frontal lobe in infancy is indeed involved in learning sequential dependency and modulates responses to deviant stimuli depending on sequential context. We also find evidence that this learned information is used to modulate activity in occipital and temporal lobes through feedback neural connections. This second finding demonstrates both the functional availability of long-range neural connections early in infancy (i.e., the connections between the frontal and occipital/temporal lobes) and that the infant brain exhibits highly plastic and dynamic responses to sensory input based on the context in which this input occurs.

## Acknowledgements

We thank all of the families who volunteered their time to participate in this study. With your participation, we now know more about how the infant brain learns and develops! We also wish to thank the research staff in the Princeton Baby Lab who provided support for this work: Carolyn Mazzei, Alex Boldin, Kachina Allen, Claire Robertson and the many wonderful undergraduate research assistants. This study was funded by the Eric and Wendy Schmidt Transformative Technology Fund at Princeton University, the Bill and Melinda Gates Foundation (INV-005792), McDonnell Foundation (AWD1005451) and the National Institutes of Health (NICHD, K99-R00 4R00HD076166-02).

## Supplementary Materials

**Figure S1.**
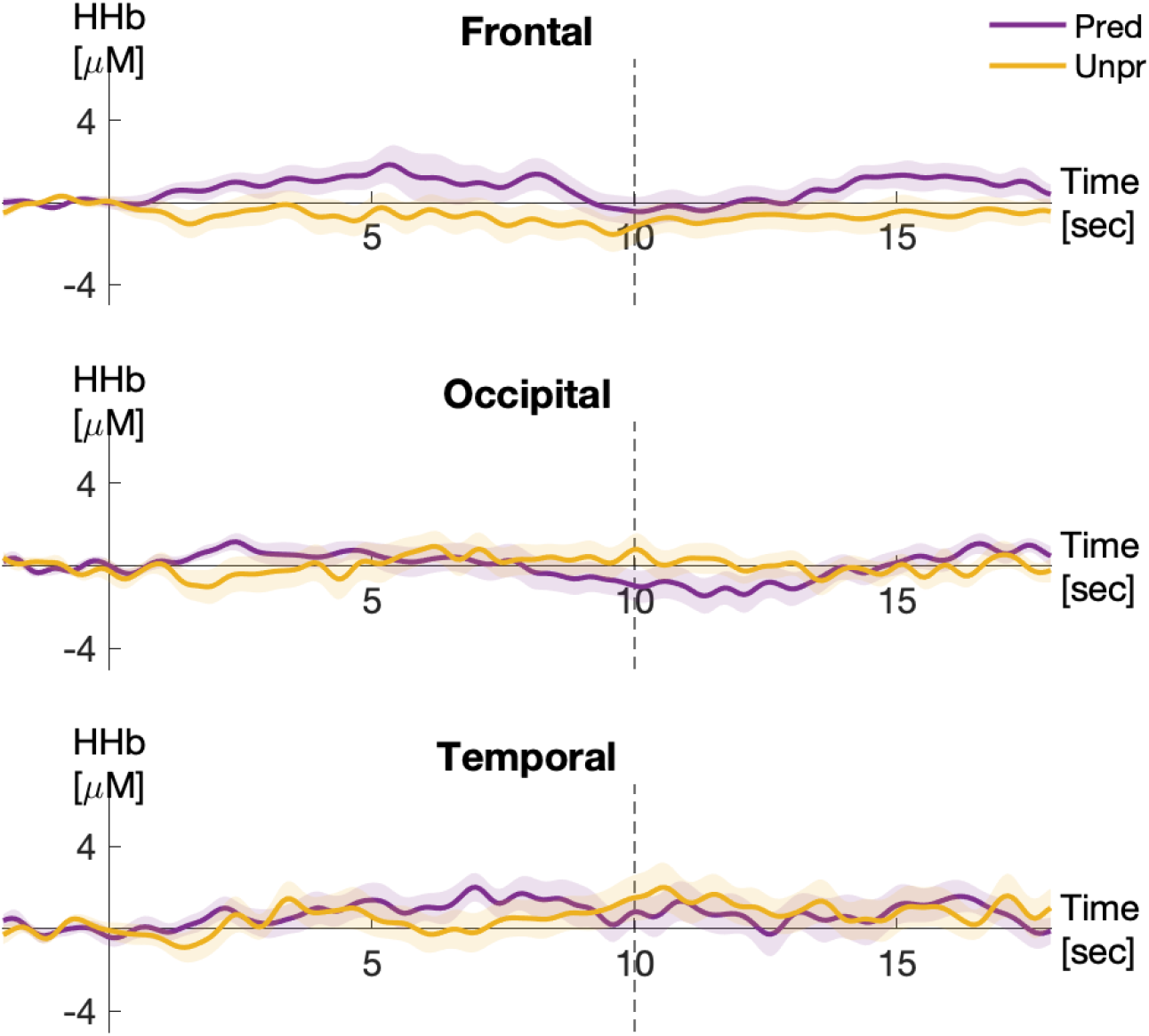
Deoxy hemoglobin concentration changes in response to the two experimental conditions.

In the main article, we report results that infants exhibit attenuated sensory responses when presented with stimuli within a predictable sequence. Convergent findings suggest that the frontal lobe is crucial for supporting learning of these sequences, while the attenuated sensory responses are in posterior sensory cortices (temporal and occipital cortices, corresponding to auditory and visual responses respectively). Thus, we hypothesized that these attenuated responses to predictable stimuli would be supported by connectivity between the frontal lobe and the posterior sensory cortices. To test this prediction, we developed a pipeline to assess task-based functional connectivity, using a background connectivity approach, with infant fNIRS data. Here, we present details on this analytic pipeline including our validation of key analytic decisions that were used to produce robust estimates of task-based connectivity. Notably however, we do not find statistically significant differences in background connectivity between the conditions examined in this study.

Background connectivity is an analytic approach to elucidating functional connectivity during task-based stimulation without the confounding effect of the temporal structure of stimulus presentation. In the presence of external stimulation, hemodynamic responses in different neural areas are correlated not only due to the connectivity between them, but also due to the synchronized task-evoked responses. If the task-evoked responses are not appropriately controlled for, the functional connectivity analysis can yield over-inflated correlation estimates (Cole et al., 2019). The background connectivity approach to inferring functional connectivity models and linearly regresses stimulus-evoked responses out of the data, before measuring correlations in the residual spontaneous fluctuations (Al-Aidroos, Said, & Turk-Browne, 2012).

Specifically, we modeled task-evoked activations by fitting a generalized linear model (GLM) using an finite impulse responses (FIR) approach (Santosa, Zhai, Fishburn, & Huppert, 2018)We chose FIR as a basis function to approximate the hemodynamic response, particularly given that we do not have definitive knowledge about the shape of the hemodynamic response in infant subjects. We defined our impulse response window as 20 seconds starting at stimulus onset to account for 10 seconds of stimulus and 10 seconds of inter-trial baseline period. At one FIR regressor per second, we used 20 FIR regressors to model the neural response, and the weighted sum of those impulses were used to estimate the shape of the hemodynamic response. We used this estimate of hemodynamic response as the measure of task-evoked cortical activity in the functional connectivity analyses.

The residuals from this model (i.e., what remains after the task-evoked responses were removed) were used to measure background activity. After confirming that the residuals were normally distributed and therefore were suitable for Pearson correlation, we calculated temporal Pearson correlations over the residuals to infer functional connectivity between the frontal, temporal and occipital lobes. The correlation coefficients (Pearson’s r) of each subject were Fisher transformed to z-values to aggregate across all subjects. Then, the z-values were Fisher transformed back to Pearson’s r values.

We first sought to evaluate the influence of some key analytic decisions in this pipeline, to determine how they affected our estimates of the background connectivity patterns. We highlight two such decisions.

### Not removing the global mean

Similar techniques have been used with fMRI data (Al-Aidroos et al., 2012), although, unlike with fMRI data, we did **not** first regress out nuisance and global variables due to the lack of coverage of non-responsive regions like brain stems and ventricles in an fNIRS recording. Indeed, we found that regressing out the global mean resulted in uniformly negative task-based connectivity patterns. In the attached figures, model 2 reflects the pipeline in which the global mean is first regressed out. This pattern of findings indicates that the regression of global or nuisance variables may not be appropriate with fNIRS data. However, in fNIRS studies including data from short-channels or which have broader coverage of non-responsive regions, it would be worthwhile to evaluate whether the inclusion of global mean or nuisance regressors in the GLM would be beneficial.

### Averaging channels before estimating background connectivity

Another analytic decision that we evaluated is when in the pipeline to average data across channels. We were interested in connectivity patterns across lobes of the brain. Since individual fNIRS channels are smaller than these lobular ROIs, and there are therefore several such channels in each lobe, we needed to average across channels to produce lobe-based connectivity patterns. Our pipeline averaged the fNIRS data into the lobular ROIs before estimating background connectivity. Our primary motivation for this decision was that we expected averaging at this early stage to increase the quality of the signals we used to estimate background connectivity. To evaluate the influence of this decision, we contrasted this to the alternative approach of calculating background connectivity for each pair of channels between two lobular ROIs and then averaging the resulting background connectivity patterns (i.e., the Fisher transformed *r* scores). As predicted, this alternative approach produced much weaker connectivity patterns. This is illustrated in Model 3 in the figures below.

After establishing this pipeline, we tested whether background connectivity was different in response to PS versus UPS, using a two-tailed *t*-test. No *t*-value was close to the threshold for statistical significance (see figures below for matrices of these *t*-values).

**Figure S2:**
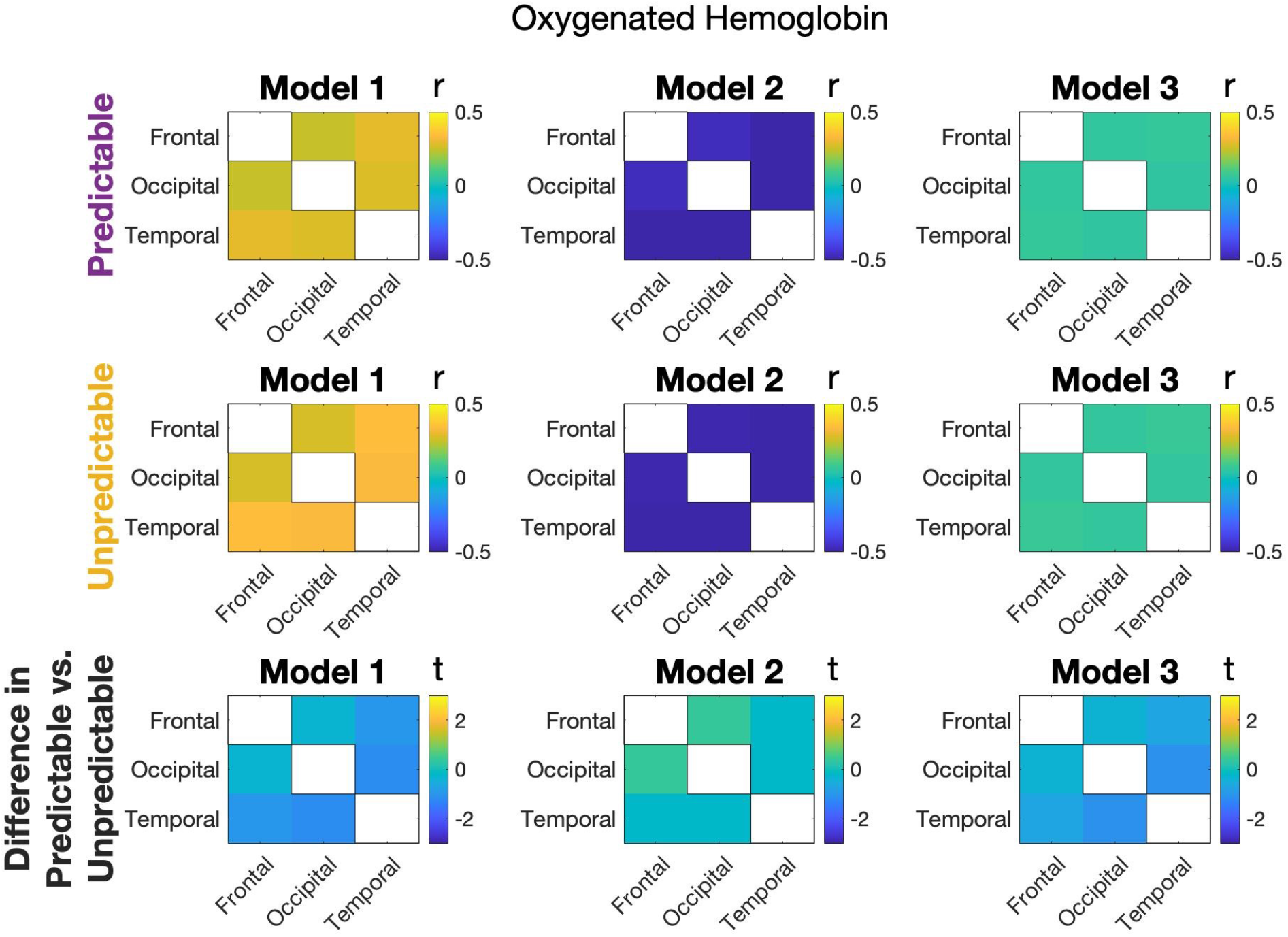
Connectivity matrices for predictable and unpredictable conditions (top and middle rows, respectively) across three models, for the oxygenated hemoglobin data. Model 1 is the established pipeline: does not remove the global mean, regresses out task-evoked activity using an FIR model, and averages responses within lobes before calculating background connectivity. Model 2: presents the same data as Model 1 but with the global mean first removed. Model 3: presents the same data as Model 1 but with the background connectivity measures estimated for all channel pairs across lobes, and then the resulting correlation coefficients averaged to produce the lobe level results. The bottom row presents the *t*-values contrasting connectivity measures between the two conditions for each model. No reliable differences were observed between conditions.

**Figure S3:**
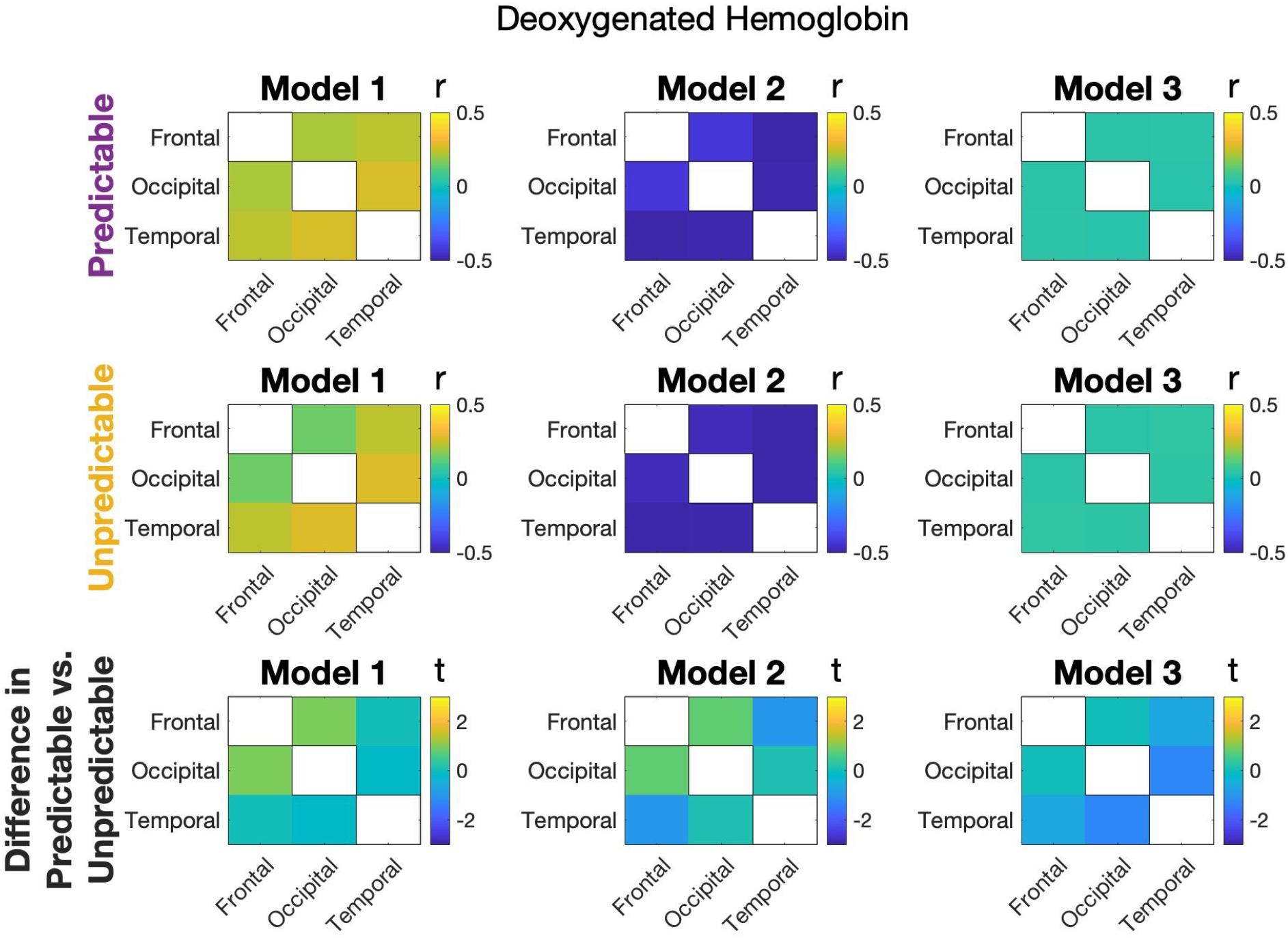
Connectivity matrices for predictable and unpredictable conditions (top and middle rows, respectively) across three models for the deoxygenated hemoglobin data. Model 1 is the established pipeline: does not remove the global mean, regresses out task-evoked activity using an FIR model, and averages responses within lobes before calculating background connectivity. Model 2: presents the same data as Model 1 but with the global mean also removed. Model 3: presents the same data as Model 1 but with the background connectivity measures estimated for all channel pairs across lobes and then the resulting correlation coefficients averaged to produce the lobe level results. The bottom row presents the t-values contrasting connectivity measures between the two conditions for each model. No reliable differences were observed between conditions.

